# Identification of DISE-inducing shRNAs by monitoring cellular responses

**DOI:** 10.1101/186890

**Authors:** Monal Patel, Marcus E. Peter

## Abstract

Off-target effects (OTE) are an undesired side effect of RNA interference (RNAi) caused by partial complementarity between the targeting siRNA and mRNAs other than the gene to be silenced. The death receptor CD95 and its ligand CD95L contain multiple sequences that when expressed as either si-or shRNAs kill cancer cells through a defined OTE that targets critical survival genes. Death induced by survival gene elimination (DISE) is characterized by specific morphological changes such as elongated cell shapes, senescence-like enlarged cells, appearance of large intracellular vesicles, release of mitochondrial ROS followed by activation of caspase-2, and induction of a necrotic form of mitotic catastrophe. Using genome-wide shRNA lethality screens with eight different cancer cell lines, we recently identified 651 genes as critical for the survival of cancer cells. To determine whether the toxic shRNAs targeting these 651 genes contained shRNAs that kill cancer cell through DISE rather than by silencing their respective target genes, we tested all shRNAs in the TRC library derived from a subset of these genes targeting tumor suppressors (TS). We now report that only by monitoring the responses of cancer cells following expression of shRNAs derived from these putative TS it was possible to identify DISE-inducing shRNAs in five of the genes. These data indicate that DISE in general is not an undefined toxic response of cells caused by a random OTE but rather a specific cellular response with shared features that points at a specific biological function involving multiple genes in the genome.

## Introduction

RNA interference is a widely used tool to reduce the expression of mRNAs. RNAi is initiated by double-stranded (ds)RNAs or pre-microRNAs, which are cleaved by Dicer, an RNase III enzyme, producing 21–23 nucleotide short interfering (si)RNAs or micro (mi)RNAs respectively, containing 2nt 3’ overhangs ^1, 2^. The antisense (guide) strand is then loaded onto the endonuclease argonaute 2 (Ago2) in the RNA-induced silencing complex (RISC) and directs the downstream targeting events mostly through complete complementarity between positions 2–8 (the seed region) at the 5′ end of the guide strand and a matching sequence (seed match) in the 3'UTR of targeted mRNAs. ^3-6^. While in case of miRNAs, the guide strand recruits the RISC to the 3’ untranslated regions (UTRs) of partially complementary mRNAs to promote translation repression or mRNA cleavage ^7, 8^, the guide strand of an siRNA is designed to be fully complementary to the target mRNA and directs the enzymatic cleavage of the mRNA by the Ago2 protein ^9-11^. RNAi can be induced by either transfecting cells with siRNAs, or by introducing short hairpin (sh)RNAs in the form of expression vectors or viruses. Apart from the intended target, the guide strands of siRNAs also recognize many mRNAs with partial complementarity in a manner similar to miRNAs, mostly involving the guide RNA seed, and studies have suggested that 3’UTR complementarity to si/shRNA seed sequences can mediate gene silencing based on an off-target effect (OTE) both through translational repression and mRNA degradation ^12-15^. In addition, improper loading of the sense/passenger strand can also lead to OTEs ^16^. This can be caused by imprecise cleavage of shRNAs by Dicer prior to RISC loading ^16^. The main causes of OTE are therefore cross-reactivities of either the guide RNA or the passenger strand loaded into the RISC ^17, 18^ with transcripts of undesired genes in the genome. The goal for virtually all RNAi projects is to selectively silence targeted genes with little or no OTE. In fact, most of the latest generation siRNAs are chemically modified to increase their stability, specificity, and to reduce OTE ^19^, and shRNAs are expressed using optimized vector systems that allow preferential loading of guide strand into the RISC ^18, 20^.

We recently reported that >80% of 22 different nonoverlapping si-, Dsi-or shRNAs derived from either CD95L or CD95 killed cancer cells by activating multiple cell death pathways ^21, 22^. Activation of the CD95/Fas surface receptor upon binding to its cognate ligand (CD95L) induces apoptosis. The CD95/CD95L system is used by immune cells to eliminate virus-infected and cancer cells through the secretion of CD95 ligand (CD95L) ^23^. Hence, the CD95/CD95L system has a tumor suppressive function. Interestingly, what at first appeared to be cancer cell death caused by silencing the expression of these two genes, actually turned out to be initiated by a mechanism completely independent of the presence of CD95 or CD95L gene products ^22^. We demonstrated that the toxic si-or shRNAs derived from either CD95 or CD95L killed cells in what appeared to be a combination of apoptosis, necrosis and mitotic catastrophe, mediated by the release of mitochondrial ROS, activation of caspase-2, and DNA damage. Morphologically, most cells responded by forming oddly shaped elongated cell structures, with likely stress induced large vesicles, and anaphase bridges ^21^. While some cells died as early as one day after introducing the toxic shRNAs, most cells died when attempting to divide ^21^. This form of cell death could not be inhibited and cancer cells had a hard time developing resistance both *in vitro* and *in vivo* ^21, 24^. We recently presented data to suggest that cells actually die through an OTE that results in the *preferential* targeting of the 3'UTRs of a set of critical survival genes ^22^. We have therefore named this form of cell death DISE (for death induced by survival gene elimination).

The discovery of DISE raised a number of puzzling questions: Why did the cancer cells appear to respond to the toxic shRNAs in a highly similar way? Why would an OTE not result in a variety of unintended cellular responses, depending on what gene or sets of genes are affected? In this study we set out to identify novel toxic shRNAs derived from a small subset of putative tumor suppressor genes other than CD95 and CD95L. Solely by monitoring cellular responses (morphology, biochemical changes, and ability to divide) by the cancer cells we have identified shRNAs derived from 5 putative tumor suppressive genes that can kill multiple cancer cells by an OTE in the absence of the coded protein that resembles DISE. We propose that these RNAi active sequences can be used to kill cancer cells.

## Results

### A subset of genes recently found to be critical for the survival of cancer cells are tumor suppressors

Previously, based on 12 shRNA-based lethality screens of 8 human cancer cell lines/cell line variants (HeLa, S3, HeLa N10, CHP-100, FU-UR-1, HEK293, A549 EGFRB, A549, H2030), we nominated 651 out of ~18,0000 genes targeted (by ~78,000 shRNAs, individually tested) as critical survival factors for cancer cells ^21^. Included were all genes for which at least 3 out of 5 shRNAs (H factor = 60) reduced cell viability more than 95% in at least 9 out of 12 independent screens ^21^. Most of the 651 genes had genuine survival functions and included genes coding for ribosomal proteins, cell cycle regulators or all three RAS genes (see Table S2 in ^21^). However, a survival function was not immediately obvious for a number of these genes and therefore they could be sources of DISE-inducing shRNAs. To increase the chance of finding such toxic shRNAs, we decided to focus on a subset of genes most unlikely to be required for cancer cell survival: tumor suppressors (TS). To identify potential TS among the 651 genes identified as survival genes, we compared the 651 genes with a curated list of 637 putative TS genes ^25^. This analysis resulted in 17 putative TS genes (plus CD95L) for which up to 94% of the targeting shRNAs killed a number of cancer cell lines in the shRNA lethality screen (**Fig. S1**). For each of the 17 genes, tumor suppressive activities have been described for various cancers (see legend of **Fig. S1B**).

### Identification of RNAi active toxic sequences derived from certain tumor suppressors

The finding that shRNAs derived from TS can kill cancer cells suggested that they may not act by reducing protein levels of their targeted genes, but by another mechanism, possibly DISE. We therefore decided to first validate the toxicity by testing five shRNAs per gene, a total of 85 shRNAs. Because we were only interested in shRNAs that killed all cancer cells, we chose three additional cell lines for this test, which were not part of the original shRNA lethality screen: HeyA8 (ovarian cancer), T89G (glioblastoma), and HCT116 (colon cancer). The latter two cell lines were chosen because we used them before to study and biochemically characterize DISE ^21^. We decided on a sequential strategy: test the shRNAs on HeyA8 cells, then test the toxic ones on T98G cells and test all shRNAs that killed these two cell lines on HCT116 cells. To identify two shRNAs per gene that killed HeyA8 cells, the 85 shRNAs (in the pLKO backbone) were screened in 96 well plates targeting the 17 TSs using a Thermo Multidrop Combi and a Tecan Freedom EVO200 for infecting cells at an MOI of five. After puromycin selection the effect on growth was monitored in the IncuCyte Zoom. For each gene, the two shRNAs that caused the strongest growth reduction were identified and used for further analysis (data not shown). This resulted in the identification of 34 toxic shRNAs targeting the 17 TS.

Because we were interested in determining if these shRNAs had similar activities and elicited cellular responses similar to the DISE-inducing shRNAs derived from CD95 or CD95L, we retested these 34 toxic shRNAs at an MOI of three on HeyA8, T98G and HCT116 cells. This was done again in the IncuCyte Zoom and growth reduction (50% reduction compared to cells infected with a nontargeting shRNA at half maximal confluency) was used as an initial surrogate marker for cell death. All 34 shRNAs targeting the 17 TS were identified as toxic to HeyA8 cells, validating the original screen. (**Fig. 1A**). Not surprisingly, while shRNAs against the tested TS were toxic, when we tested 4-5 shRNAs targeting two of the most widely studied and most highly mutated TS in human cancers, p53 and PTEN, none of them qualified as toxic shRNAs using the threshold we had defined (**Fig. 1B**). This suggested that only certain TS contain RNAi active sequences that can kill cancer cells. Of the 34 shRNAs targeting the 17 TS genes, 30 shRNAs were also toxic to T98G cells targeting 15 of the TSs (**Fig. S2A**). Because we were only interested in shRNAs that kill all three cancer cells, we only tested these 30 shRNAs on HCT116 cells (**Table 1** and **Fig. S2B**). This reduced the number of shRNAs that killed all three cell lines to 26 shRNAs (targeting 13 TS).

**Figure 1.**
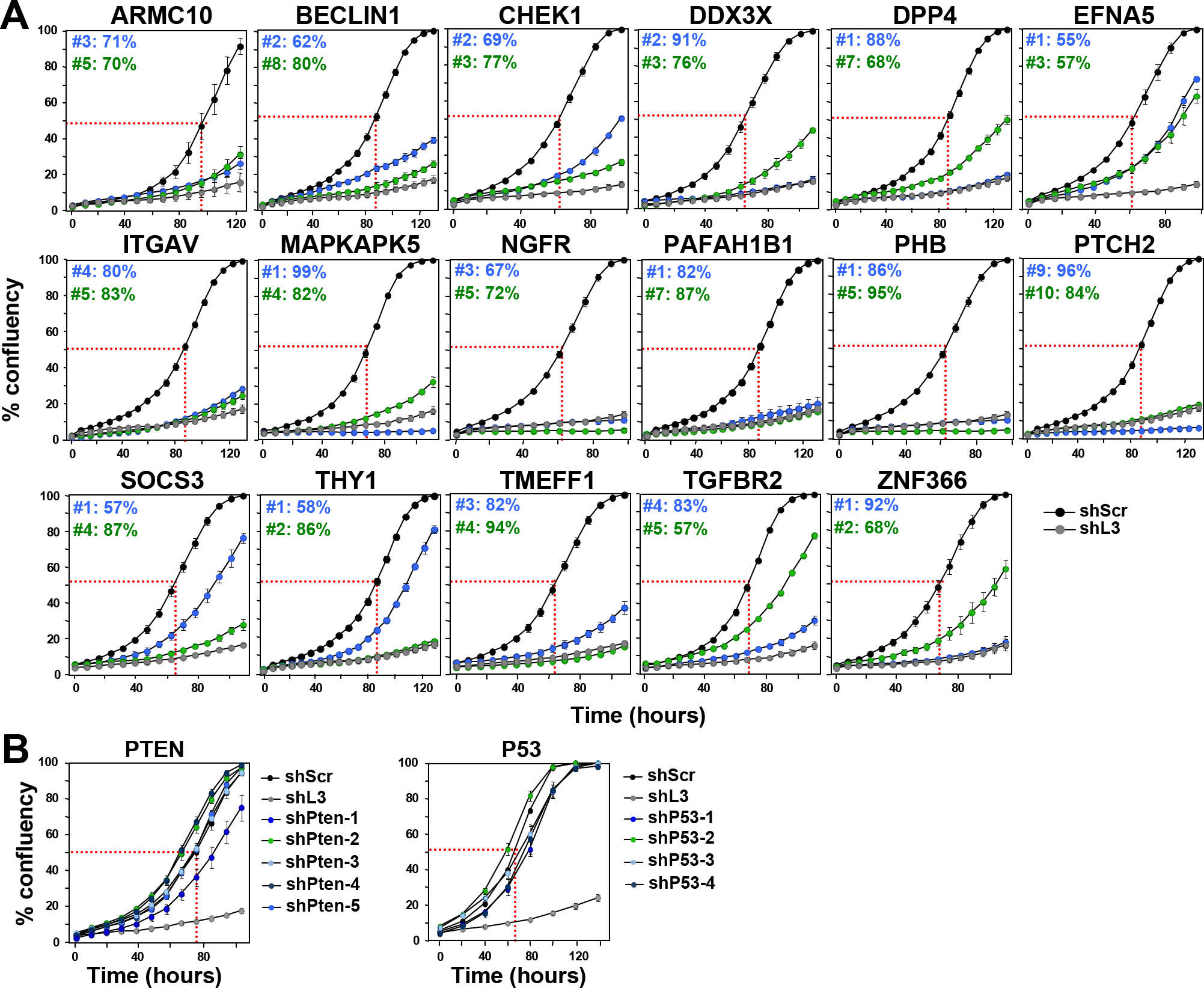
shRNAs derived from 17 TS genes cause growth reduction in HeyA8 cells. (A) Percent cell confluence over time of HeyA8 cells after infection with shScr, shL3 and two shRNAs for each of the 17 TS genes. The curves for cells infected with two independent shRNAs for each TS gene and their specific ID number and respective growth reduction caused by each shRNA are shown in blue and green. Percent growth reduction values (as shown in Table 1) were calculated using STATA1C software when cells infected with shScr reached half maximal confluency as indicated by the red dotted line. (B) Percent cell confluence over time of HeyA8 cells after infection with shScr, shL3, and five shRNAs targeting PTEN (left) and four shRNAs targeting p53 (right). Percent growth reduction values are shown in **Table 1**.

**Table 1:** Summary of all assays leading to the identification of DISE inducing TS-derived shRNAs. Results of systematic analyses (left to right) of two shRNAs to each of the 17 TS in various assays.

| TS | shRNA ID# | TRC# | % growth reduction compared to shScr |  |  | Morphological changes similar to DISE (in both HeyA8 & T98G) | Caspase 2 activity (in HeyA8 cells) | ROS (in HeyA8 cells) | % growth reduction in HAP1 cells | % growth reduction in respective k.o. cells |  |
| --- | --- | --- | --- | --- | --- | --- | --- | --- | --- | --- | --- |
|  |  |  | HeyA8 | T98G | HCT116 |  |  |  |  |  |  |
| PAFAH1B1 | #1 | TRCN0000050966 | 82 | 57 | 78 | yes | yes | yes | 0 |  |  |
|  | #7 | TRCN0000050964 | 87 | 90 | 88 | yes | yes | yes | 29 |  |  |
| NGFR | #3 | TRCN0000058155 | 67 | 72 | 27 |  |  |  |  |  |  |
|  | #5 | TRCN0000058157 | 72 | 51 | 86 |  |  |  |  |  |  |
| ITGAV | #4 | TRCN0000010769 | 80 | 99 | 91 | no |  |  |  |  |  |
|  | #5 | TRCN0000010768 | 83 | 80 | 85 | no |  |  |  |  |  |
| DPP4 | #1 | TRCN0000050773 | 88 | 87 | 78 | yes | yes | yes |  |  |  |
|  | #7 | TRCN0000050776 | 68 | 58 | 73 | yes | no | no |  |  |  |
| EFNA5 | #1 | TRCN0000058218 | 55 | 63 |  |  |  |  |  |  |  |
|  | #3 | TRCN0000058220 | 57 | 0 |  |  |  |  |  |  |  |
| TMEFF1 | #3 | TRCN0000073510 | 82 | 88 | 89 | yes | yes | yes | 90 | 89 | TMEFF1 |
|  | #4 | TRCN0000073511 | 94 | 99 | 91 | yes | yes | yes | 93 | 85 |  |
| CHEK1 | #2 | TRCN0000009947 | 69 | 76 | 88 | no |  |  |  |  |  |
|  | #3 | TRCN0000039856 | 77 | 86 | 86 | yes |  |  |  |  |  |
| PTCH2 | #9 | TRCN0000033327 | 96 | 96 | 94 | yes | yes | yes | 90 |  |  |
|  | #10 | TRCN0000033328 | 84 | 83 | 84 | yes | yes | yes | 0 |  |  |
| ARMC10 | #3 | TRCN0000130777 | 71 | 95 | 88 | yes | yes | yes | 60 | 93 | ARMC10 |
|  | #5 | TRCN0000128466 | 70 | 86 | 57 | yes | yes | yes | 73 | 95 |  |
| BECN1 | #2 | TRCN0000033552 | 62 | 39 |  |  |  |  |  |  |  |
|  | #8 | TRCN0000033553 | 80 | 97 |  |  |  |  |  |  |  |
| DDX3X | #2 | TRCN0000000002 | 91 | 80 | 96 | yes |  |  |  |  |  |
|  | #3 | TRCN0000000003 | 76 | 92 | 64 | no |  |  |  |  |  |
| THY1 | #1 | TRCN0000057023 | 58 | 60 | 83 | no |  |  |  |  |  |
|  | #2 | TRCN0000057024 | 86 | 75 | 69 | yes |  |  |  |  |  |
| PHB | #1 | TRCN0000029204 | 86 | 89 | 99 | no |  |  |  |  |  |
|  | #5 | TRCN0000029208 | 95 | 91 | 98 | no |  |  |  |  |  |
| SOCS3 | #1 | TRCN0000057073 | 57 | 72 | 55 | yes | yes | yes | 90 | 93 | SOCS3 |
|  | #4 | TRCN0000057076 | 87 | 97 | 88 | yes | yes | yes | 85 | 78 |  |
| ZNF366 | #1 | TRCN0000020134 | 92 | 93 | 46 |  |  |  |  |  |  |
|  | #2 | TRCN0000020135 | 68 | 58 | 56 |  |  |  |  |  |  |
| MAPKAPK5 | #1 | TRCN0000000681 | 99 | 98 | 94 | yes | yes | yes | 97 | 98 | MAPKAPK5 |
|  | #4 | TRCN0000195129 | 82 | 65 | 71 | yes | yes | yes | 88 | 78 |  |
| TGFB2 | #4 | TRCN0000195606 | 82 | 90 | 83 | yes | yes | yes | 91 | 77 | TGFB2 |
|  | #5 | TRCN0000197056 | 57 | 84 | 62 | yes | yes | yes | 95 | 71 |  |
| PTEN | #1 | TRCN0000355840 | 0 |  |  |  |  |  |  |  |  |
|  | #2 | TRCN0000355841 | 0 |  |  |  |  |  |  |  |  |
|  | #3 | TRCN0000355842 | 0 |  |  |  |  |  |  |  |  |
|  | #4 | TRCN0000355843 | 0 |  |  |  |  |  |  |  |  |
|  | #5 | TRCN0000355946 | 29 |  |  |  |  |  |  |  |  |
| P53 | #1 | TRCN0000342334 | 30 |  |  |  |  |  |  |  |  |
|  | #2 | TRCN0000342335 | 0 |  |  |  |  |  |  |  |  |
|  | #3 | TRCN0000003754 | 19 |  |  |  |  |  |  |  |  |
|  | #4 | TRCN0000342259 | 22 |  |  |  |  |  |  |  |  |

shRNAs were effective
shRNAs were not effective
shRNAs were effective shRNAs were not effective

### Toxic TS derived shRNAs trigger DISE-like cell death

To determine which of the shRNAs killed the three cell lines in a fashion similar to DISE induction observed with CD95L derived shRNAs, we compared the morphological changes seen in HeyA8 and T98G cells after infection with a lentiviral shRNA with that seen in cells infected with the CD95L derived shL3. We were unable to perform a morphological analysis in HCT116 cells since the cells were too small and without clear morphological features to distinguish. In HeyA8 cells, 3-7 days after infection, we detected the typical stress-induced elongated cell shapes (**Fig. 2A**), appearance of large intracellular vesicles, and enlargement and senescence-like cell flattening (**Fig. 2B**). 8 of the 13 remaining TS genes had shRNAs that both elicited these changes (**Fig. 2** and **Table 1**). In T98G cells, for the same 8 genes (all of the 16 shRNAs), we observed that the cells attempted to divide and immediately after that they rounded up and died (data not shown).

**Figure 2.**
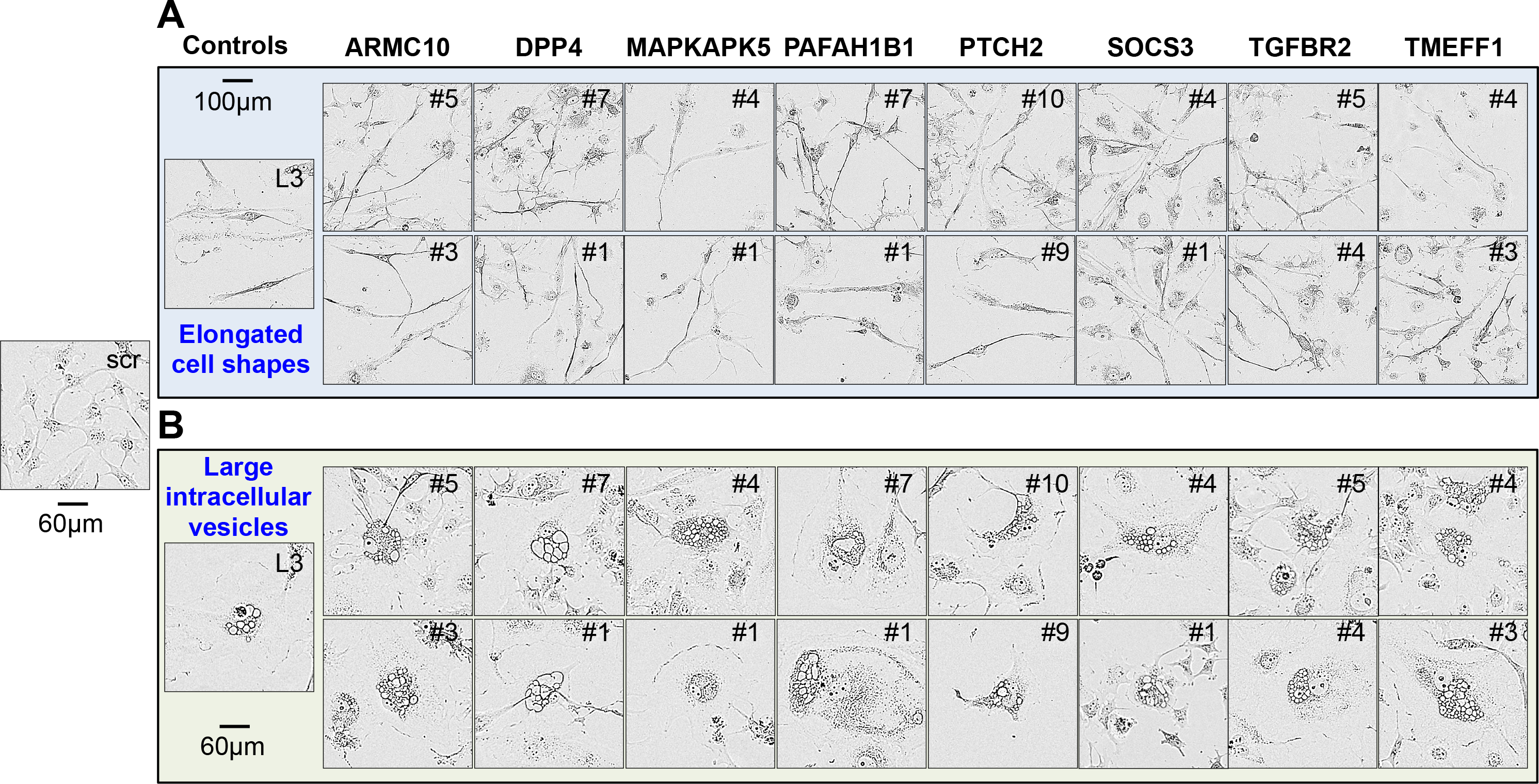
Toxic shRNAs derived from eight TS genes induce DISE-like morphological changes in HeyA8 cells. Representative phase-contrast images showing elongated cell shapes (A); enlarged, flattened cells and presence of intracellular granules in HeyA8 cells infected with shRNAs against eight of the 17 TS and shL3 (B). shScr treated cells are shown as control.

In order to determine whether the remaining 16 shRNAs induced cell death biochemically similar to DISE, we tested whether these shRNAs caused ROS production and caspase-2 activation in HeyA8 cells - two characteristic features of DISE seen in HeyA8 cells after infection with shL3 and also observed in multiple other cell lines after introducing multiple CD95L or CD95 targeting shRNAs ^21^. For 7 of the remaining TS, both shRNAs caused significant induction of ROS and caspase-2 activation (**Fig. 3A** and **B**). Our sequential analysis in three cancer cell lines allowed us to narrow down the list of potential shRNAs that killed cancer cells by DISE to seven (**Table 1**). To confirm that all shRNAs derived from these seven genes did not just result in growth reduction but actually killed cancer cells, we quantified DNA fragmentation in HeyA8 cells 8 days after lentiviral infection (**Fig. 3C**). Indeed, all 14 shRNAs caused a significant increase in subG1 DNA, suggesting that they all at various levels killed cancer cells.

**Figure 3.**
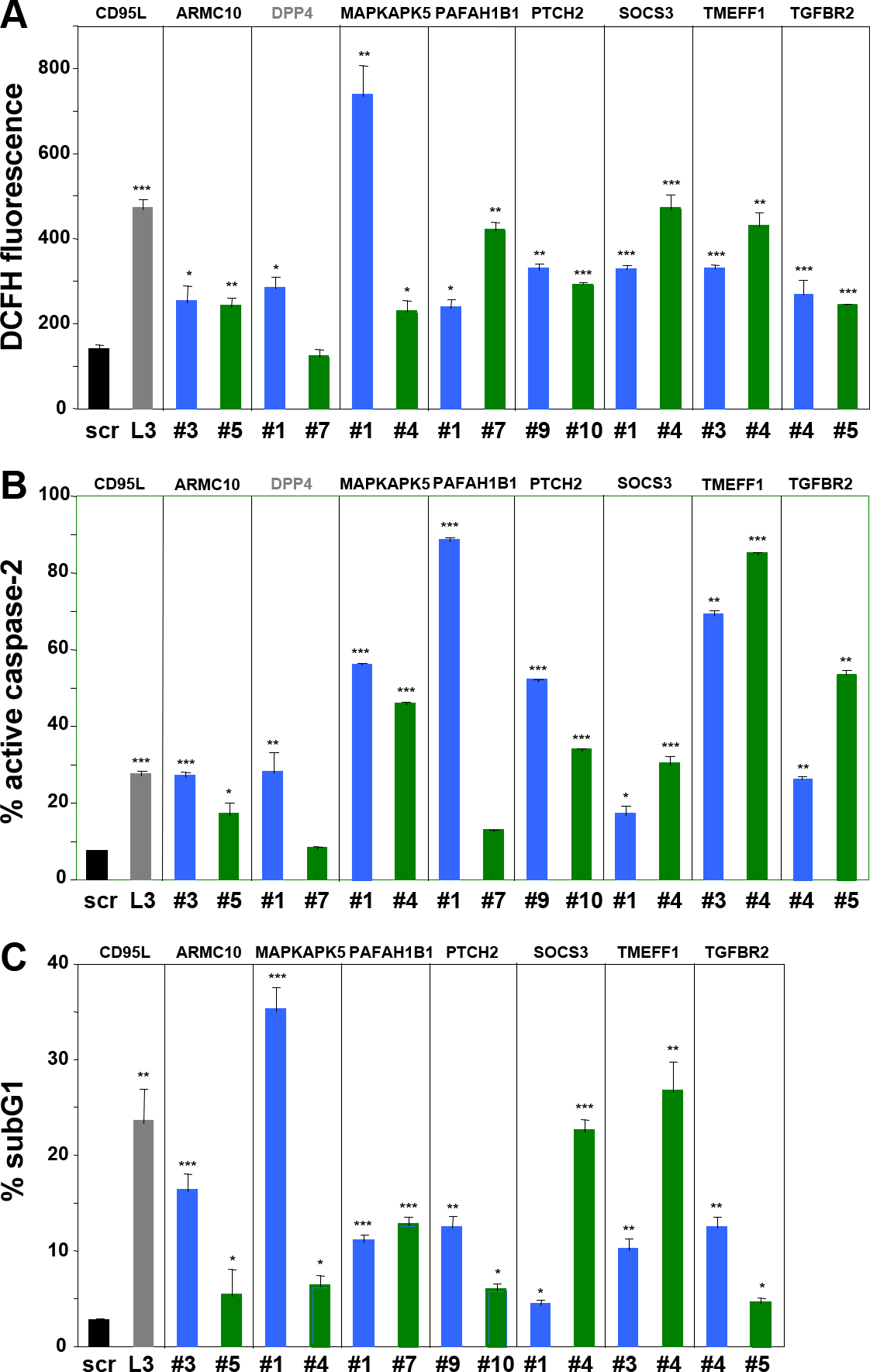
Toxic shRNAs derived from seven TS genes induce death that is biochemically similar to DISE. Quantification of ROS production by dichlorofluorescin (DCFH) fluorescence (A), caspase-2 activity (B) and quantification of cell death with PI staining (C) in HeyA8 cells 8 days after infection with shScr, shL3, and shRNAs derived from eight of the TS genes. ID numbers are shown in **Table 1**. p-values were calculated using students t-test. *p<0.01, **p<0.001, ***p<0.0001. For genes in black, both shRNAs were functionally active whereas genes shown in grey, only one the two shRNAs had a significant effect. Color code for the two shRNAs per gene is the same as in Fig. 1.

### Toxic shRNAs derived from five TS genes kill cancer cells through DISE

One of the most surprising properties of DISE-inducing shRNAs is that they kill cancer cells independent of targeting the mRNA they were designed to silence; instead we reported that these sequences are toxic to cells through a unique form of OTE that targets a network of critical survival genes ^22^. We narrowed down our list of toxic shRNAs to only include shRNAs that killed cancer cells in a way that was similar to DISE in morphology and biochemistry; we now considered whether these TS genes were enriched in shRNAs that kill cancer cells in a way independent of the expression of the coding protein, which would be indicative of the toxic OTE that is DISE. We chose to study this using HAP1 cells for two reasons: 1) They are available as knock-out cells (generated by using CRISPR/Cas9 gene editing) for most human genes (that are not essential for cell survival) and 2) We recently demonstrated that DISE-inducing shRNAs derived from either CD95 or CD95L could still kill CD95 or CD95L deficient HAP1 cells (data not shown). For five of the seven genes, both shRNAs reduced growth of unmodified HAP1 cells >50% (**Table 1, Fig. S2C**); hence these five genes could be tested in HAP1 CRISPR/Cas9 modified cells. In all HAP1 cells using CRISPR/Cas9 gene editing a frame shift mutation was introduced downstream of the translational start codon. Two of the mutant clones, ARMC10 and MAPKAPK5, were validated by Western blotting to be protein knock outs (**Fig. 4B**, far right panel). Two of the genes, SOCS3 and TMEFF1, are not expressed in HAP1 cells (Transcripts Per Kilobase Million (TPM) of less than 3 in RNA Seq analysis are considered undetectable ^26^). All of the 5 CRISPR/Cas9 modified cell lines still died after the introduction of shRNAs derived from these genes (**Fig. 4A** and **4B**, **Table 1**). Because for two of the genes the Western blot confirmation of a complete knockout was inconclusive (MAPKAPK5-multiple bands; and TGFBR2 - no band), an additional k.o clone was generated and tested. For MAPKAPK5 an out-of-frame deletion was introduced into exon 8 and for TGFBR2 in exon 4 (data not shown). Both clones were as sensitive to the two toxic shRNAs derived from these genes as wt cells (data not shown). The data indicate that all of the toxic shRNAs we identified derived from the five TS killed the cells through an OTE. Because the result of this OTE is cell death and because this cell death in all tested cell lines resembled DISE we conclude that these genes contain toxic sequences that can kill cancer cells by DISE. These data suggest that CD95 and CD95L are not unique and that the human genome likely contains multiple genes that contain sequences that have DISE inducing activities when expressed as small double stranded RNAs.

**Figure 4.**
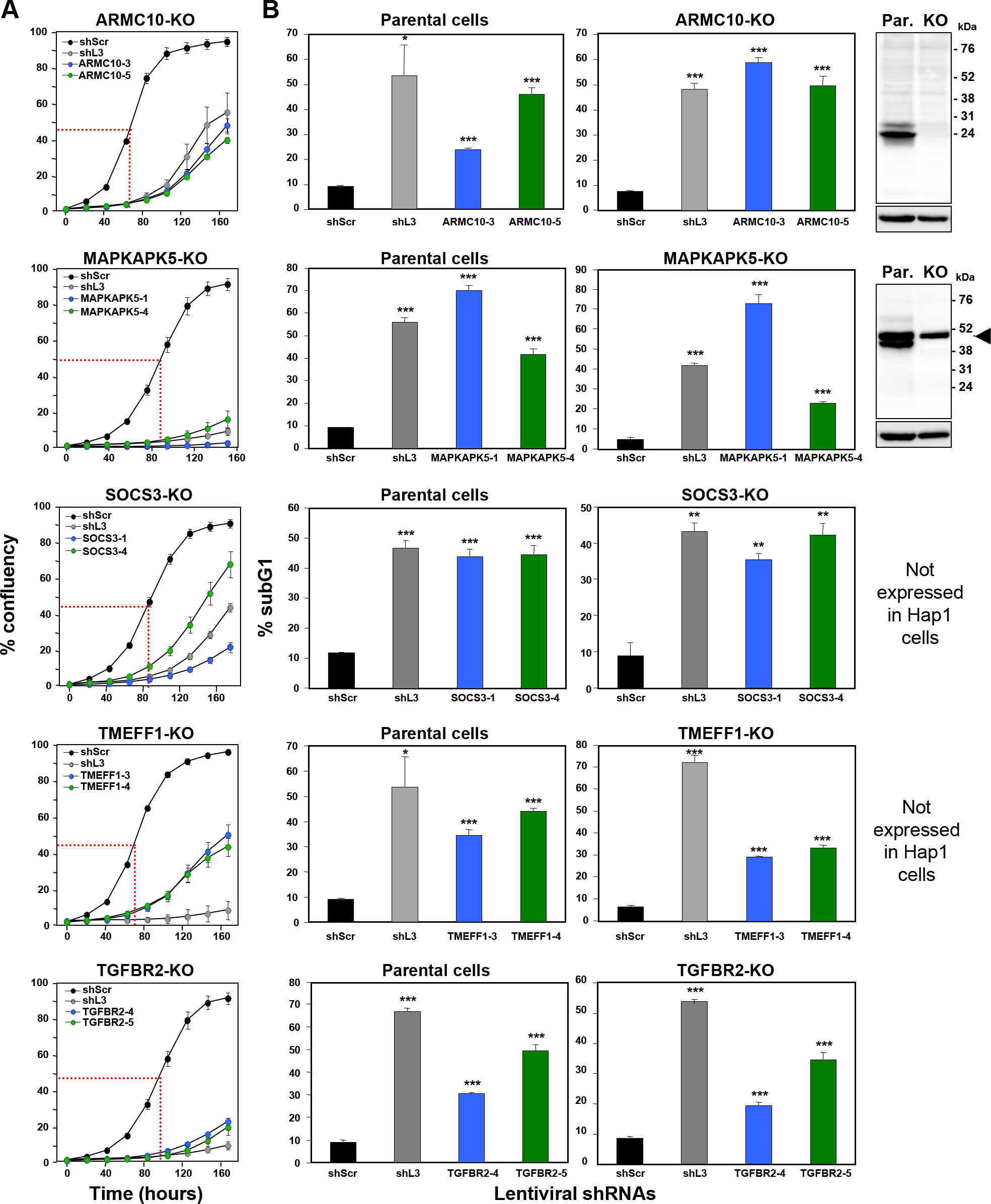
shRNAs derived from four TS genes kill cells in the absence of the transcript and/or protein produced from the targeted gene. Percent cell confluence over time (A) and percent nuclear PI staining (B) of HAP1 parental cells, or HAP1 knock-out cells, after infection with shScr, shL3, or one of two shRNAs each targeting the respective TS. Percent growth reduction values (as shown in Table 1) were calculated using STATA1C software when cells infected with shScr reached half maximal confluency as indicated by the red dotted line. p-values were calculated using a t-test. Western blot analyses in (B) confirm the knock-out of ARMC10 and MAPKAPK5 at the protein level. Arrowhead marks likely unspecific band. *p<0.01, **p<0.001, ***p<0.0001. Color code for the two shRNAs per gene is the same as in Fig. 1.

## Discussion

RNAi has become one of the most utilized methods to study the function of genes. Countless reports of gene-specific silencing and genome-wide screens document the power of RNAi ^21, 27-30^. However, one caveat of RNAi screens is the OTE. This can be caused by cross-reactivities between the guide strand of the siRNA and the target mRNAs due to partial complementarity. In addition, OTE can be caused by unintended loading of the passenger strand into the RISC. OTE has been described to occur ^16^ and was found to affect many genes and to only require a complementarity of 6-7 nucleotides between the targeting si/shRNA and the affected mRNAs ^12-14^. However, studies were unable to predict which genes would be and which genes would not be affected by OTE ^13^. This finding is consistent with the assumption that OTEs are truly random. When all OTEs are truly random, one would expect cells to respond in various ways depending on the mRNAs affected by the OTE. Similarly, when OTEs lead to cell death, one would assume that different forms of cell death with different morphologies and different signaling pathways would be activated.

We recently identified a general OTE that results in the death of most tested cancer cells. It was found to preferentially affect transformed cells ^21^, and among them cancer stem cells ^31^. Interestingly, this OTE preferentially affected genes that are critical for cancer cell survival. We named this from of cell death DISE (death induced by survival gene elimination). Cells dying by DISE, in most cases, display similar morphologies and share a number of biochemical responses suggesting that DISE is not a random occurrence but has an underlying specific biological purpose which we are currently studying.

DISE was discovered by testing a large number of si‐ and shRNAs derived from either CD95 or CD95L. Since it was a sequence-specific OTE, it was likely that other genes also contained sequences that when expressed as shRNAs would induce DISE. We have now confirmed that shRNAs derived from a number of TS can induce a form of cell death that resembles DISE, with the same morphology, elongated cell shapes, ROS production, activation of caspase-2, inability to properly divide followed by DNA degradation and cell death. Solely by scoring similarities between the responses of cells to different toxic shRNAs did we identify 10 shRNAs (targeting 5 TS) that all killed cancer cells in which the gene was either disabled by CRISPR/Cas9 gene editing (protein knockout confirmed for two of them) or not expressed. In contrast, none of the shRNAs designed to silence the two most highly studied TS, p53 and PTEN caused significant cell death, suggesting that it is a selective group of genes that contain toxic RNAi active sequences.

Our data do not allow us to conclude that shRNAs derived from TS are particularly prone to inducing DISE. TS were merely chosen as a group of genes that were most unlikely to be critical for the survival of cancer cells. Hence, the shRNAs derived from our lethality screens designed to target TS were expected to be enriched in shRNAs that induce cell death by an OTE. Our data now suggest that certain TS-derived shRNAs can kill cancer cells through DISE and in five cases, we provide evidence to suggest that these shRNAs killed the cells in the absence of functional protein consistent with the action of the DISE mechanism. Based on these data, we propose that DISE is a general mechanism through which toxic shRNAs derived from multiple genes kill cancer cells.

## Materials and methods

### Reagents

Propidium iodide (#P4864) and puromycin (#P9620) were purchased from Sigma-Aldrich. Antibodies used for western blot were: anti-MAPKAPK5 (D70A10) rabbit mAb from Cell Signaling (#7419); anti-ARMC10 (#NBP1-81127) rabbit pAb from Novus Biologicals; and goat anti-rabbit IgG human adsorbed-HRP #4010-05) was from Southern Biotech.

### Cell lines

The ovarian cancer cell line HeyA8, and the colon cancer cell line HCT116 were grown in RPMI 1640 medium (Mediatech Inc), supplemented with 10% heat-inactivated FBS (Sigma-Aldrich), 1% L-glutamine (Mediatech Inc), and 1% penicillin/streptomycin (Mediatech Inc). The glioblastoma cell line T98G was grown in EMEM (ATCC#30-2003), containing 10% heat-inactivated FBS, 1% L-Glutamine, and 1% penicillin/streptomycin. The chronic myelogenous leukemia cell line HAP1 (Horizon Discovery #C631), HAP1 ARMC10 k.o. (Horizon Discovery # HZGH005198c009, 2 bp deletion in exon 2, k.o. validated by Western blotting), HAP1 TGFBR2 k.o. (Horizon Discovery # HZGHC000035c015, 13 bp deletion in exon 1, and cat# HZGHC006289c002 - 7 bp deletion in exon 4, protein not detectable by Western blotting), HAP1 TMEFF1 k.o. (Horizon Discovery # HZGHC005199c011, 2 bp deletion in exon 2, k.o. not validated by Western blotting), HAP1 SOCS3 k.o. (Horizon Discovery # HZGHC005447c010, 25 bp deletion in exon 2, protein not detectable by Western blotting), and HAP1 MAPKAPK5 k.o. (Horizon Discovery # HZGHC000217c004, 4 bp deletion in exon 2, and cat# HZGHC006287c012 - 4 bp deletion in exon 8, k.o. validated by Western blotting) cell lines, were cultured in Gibco IMDM (Life Technologies #12440053), supplemented with 10% heat-inactivated FBS, 1% L-Glutamine, and 1% penicillin/ streptomycin.

### Knockdown via lentiviral shRNAs

Cells were infected with the following MISSION^®^ Lentiviral Transduction Particles (Sigma): pLKO.1-puro Control Transduction Particle coding for a nontargeting (scrambled) shRNA (#SHC002V), shRNAs against mRNA NM_000430 (Homo sapiens *PAFAH1B1*) TRCN0000050966 (#1: TGACCATTAAACTATGGGATT) and TRCN0000050964 (#7: CGTATGGGATTACAAGAACAA), shRNAs against mRNA NM_002507 (Homo sapiens *NGFR*) TRCN00000058155 (#3: CCTCCAGAACAAGACCTCATA) and TRCN00000058157 (#5: GCCTACGGCTACTACCAGGAT), shRNAs against mRNA NM_002210 (Homo sapiens *ITGAV*) TRCN0000010768 (#4: GTGAGGTCGAAACAGGATAAA) and TRCN0000010769 (#5: CGACAGGCTCACATTCTACTT), shRNAs against mRNA NM_001935 (Homo sapiens *DPP4*) TRCN0000050773 (#1: GCCCAATTTAACGACACAGAA) and TRCN0000050776 (#7: GACTGAAGTTATACTCCTTAA), shRNAs against mRNA NM_010109 (Homo sapiens *EFNA5*) TRCN0000058218 (#1: GAGACCAACAAATAGCTGTAT) and TRCN0000058220 (#3: CGCGGCACAAACACCAAGGAT), shRNAs against mRNA NM_003692 (Homo sapiens *TMEFF1*) TRCN0000073510 (#3: CATGCCAATTTCAGTGCCATA) and TRCN0000073511 (#4: GCCAATTTCAGTGCCATACAA), shRNAs against mRNA NM_001274 (Homo sapiens *CHEK1*) TRCN0000009947 (#2: GACAGAATAGAGCCAGACATA) and TRCN0000039856 (#3: GCCCACATGTCCTGATCATAT), shRNAs against mRNA NM_003738 (Homo sapiens *PTCH2*) TRCN0000033327 (#9: GCTGCATTACACCAAGGAGAA) and TRCN0000033328 (#10: CGTACTCACATCCATCAACAA), shRNAs against mRNA NM_031905 (Homo sapiens *ARMC10*) TRCN0000130777 (#3: GCACATGCTTCACAGTTACAT) and TRCN0000128466 (#5: GCTTTAGTTGATCACCATGAT), shRNAs against mRNA NM_003766 (Homo sapiens *BECN1*) TRCN0000033552 (#2: CTCAAGTTCATGCTGACGAAT) and TRCN0000033553 (#8: GCTTGGGTGTCCTCACAATTT), shRNAs against mRNA NM_001356 (Homo sapiens *DDX3X*) TRCN0000000002 (#2: CGGAGTGATTACGATGGCATT) and TRCN0000000003 (#3: CGTAGAATAGTCGAACAAGAT), shRNAs against mRNA NM_006288 (Homo sapiens *THY1*) TRCN0000057023 (#1: GCCATGAGAATACCAGCAGTT) and TRCN0000057024 (#2: CGAACCAACTTCACCAGCAAA), shRNAs against mRNA NM_002634 (Homo sapiens *PHB*) TRCN0000029204 (#1: CCCAGAAATCACTGTGAAATT) and TRCN0000029208 (#5: GAGTTCACAGAAGCGGTGGAA), shRNAs against mRNA NM_003955 (Homo sapiens *SOCS3*) TRCN0000057073 (#1: CCACCTGGACTCCTATGAGAA) and TRCN0000057076 (#4: CGGCTTCTACTGGAGCGCAGT), shRNAs against mRNA NM_152625 (Homo sapiens *ZNF366*) TRCN0000020134 (#1: AGGCAGTTCAAATATAGCTTT) and TRCN0000020135 (#2: GCCCACAAAGATGCCCTATAA), shRNAs against mRNA NM_003668 (Home sapiens *MAPKAPK5*) TRCN0000000681 (#1: GCGGCACTGTCACTTGTTAAA) and TRCN0000195129 (#4: CAGTATCAATTGGACTCAGAA), shRNAs against mRNA NM_003242 (Homo sapiens *TGFBR2*) TRCN0000195606 (#4: CGACATGATAGTCACTGACAA) and TRCN0000197056 (#5: GACCTCAAGAGCTCCAATATC), shRNA targeting mRNA NM_000639 (Homo sapiens *FasLG*) TRCN0000059000 (shL3: ACTGGGCTGTACTTTGTATAT), 5 shRNAs targeting mRNA NM_000314 (Homo sapiens *PTEN*) TRCN0000355840 (#1: GGCACAAGAGGCCCTAGATTT), TRCN0000355841 (#2: ACAGTAGAGGAGCCGTCAAAT), TRCN0000355842 (#3: GACTTAGACTTGACCTATATT), TRCN0000355843 (#4: GACGAACTGGTGTAATGATAT), TRCN0000355946 (#5: ACATTATGACACCGCCAAATT), 4 shRNAs targeting mRNA NM_000546 (Homo sapiens *TP53*) TRCN0000342334 (#1: CACCATCCACTACAACTACAT), TRCN0000342335 (#2: CGGCGCACAGAGGAAGAGAAT), TRCN0000003754 (#3: TCAGACCTATGGAAACTACTT), TRCN0000342259 (#4: GTCCAGATGAAGCTCCCAGAA).

Infection was performed according to the manufacturer’s protocol. Briefly, 50,000 cells seeded the day before on a 6-well plate were infected with each lentivirus at an MOI of 3 in presence of 8μg/mL polybrene overnight. Media was changed the next day, followed by selection with 3μg/mL puromycin 24 hours later. Cells were selected for at least 48 hours, then seeded on a 96-well plate and placed in the IncuCyte (Essen Bioscience) to measure confluence or expanded for 4 days to assess cell viability with propidium iodide staining.

### ROS measurement

Intracellular ROS production was measured after 8 days of infection with lentiviral shRNAs by incubating cells with 10 μM CM-H2DCFDA (C6827; Invitrogen Molecular Probes) in media at 37°C for 30 min. CM-H2DCFDA, a cell–permeable fluorogenic probe, is cleaved by intracellular esterases forming DCFH, which in presence of ROS, gets oxidized to the fluorescent compound DCF. Following incubation, cells were washed three times with PBS, and ROS was quantified by flow cytometry.

### Caspase-2 activity measurement

Intracellular caspase-2 activity was detected *in situ* using FAM-VDVAD-FMK (ImmunoChemistry Technologies, LLC) according to the manufacturer’s instructions. Briefly, cells were harvested 8 days after infection with lentiviral shRNAs. The pellet was resuspended in 290 μl of medium, to which 10 μl of 30x FAM-VDVAD-FMK was added. Cells were incubated at 37°C for 1 hour, washed with PBS, and resuspended in 300 μl of medium. Cells were kept on ice protected from light and immediately analyzed by flow cytometry.

### Cell death assay (propidium iodide staining)

Cells infected with lentiviral shRNAs were plated in triplicates on 12 well plates after 2 days of puromycin selection, and plates were incubated at 37°C for 4 days. The total cell pellet consisting of live and dead cells was resuspended in Nicoletti buffer (0.1% sodium citrate, pH 7.4, 0.05% Triton X-100, 50 μg/ml propidium iodide). After incubating for 2-4 hours in the dark at 4°C, percent cell death was quantified by flow cytometry.

### Western blot analysis

Cells were lysed using RIPA lysis buffer (1% SDS, 1% Triton X-100, 1% deoxycholic acid) and protein concentration was determined using the DC Protein Assay kit (Bio-Rad). Equal amounts of protein (30 μg) were resolved on 10% SDS-PAGE gels and transferred to nitrocellulose membrane (Amersham Protran 0.45 μm, GE Healthcare Life Science). The membranes were blocked with 5% non-fat dry milk in 0.1% Tween-20/TBS and then incubated in primary antibodies at 4°C overnight. After washing 3 times with TBST, membranes were incubated with secondary antibodies followed by washing again. Detection was performed using the ECLTM Western Blotting Detection Reagents reagent (GE Healthcare) and developed using a chemiluminescence imager, G:BOX Chemi XT4 (Syngene). Both primary and secondary antibodies were diluted in the blocking buffer (5% milk in 0.1% Tween-20/TBS) as follows: anti-ARMC10 (1: 250), anti-MAPKAPK5 (1:1000) and goat anti-rabbit IgG human adsorbed-HRP (1.5000)

### Statistical analyses

Growth reduction was scored as significant when cell growth was inhibited at least 50% at the half maximal growth of shScr infected cells. Percent growth reduction values were calculated using the formula: [(y_1_-c_1_)-(y_2_-c_2_)]/[(y_1_-c_1_)]*100 where y_1_ is the half maximal confluency for cells infected with shScr (i.e. if the cells grew from 5% to 100% then y_1_=[(100+5)/2]); c_1_ is the starting confluency for cells infected with shScr. STATA1C software was then used to obtain the time (t_1_) for y_1_ and also to obtain the value of y_2_, which is the confluency of cells infected with TS shRNAs at t_1_, and c_2_ is their starting confluency. Experiments were performed in triplicates and the data were expressed as mean ± SD. Statistical analysis was performed using Student’s two-tailed t-test. A value of p<0.05 was considered to be significant.

## Acknowledgments

We are grateful to Denise Scholtens for biostatistics support and to Sam Bettis at the Cellular Screening Center at the University of Chicago.

## Author contributions

M.P. performed the experiments and M.E.P. designed the experiments and wrote the manuscript. Both authors read and approved the final manuscript.

## Supplementary data

**Figure S1:**
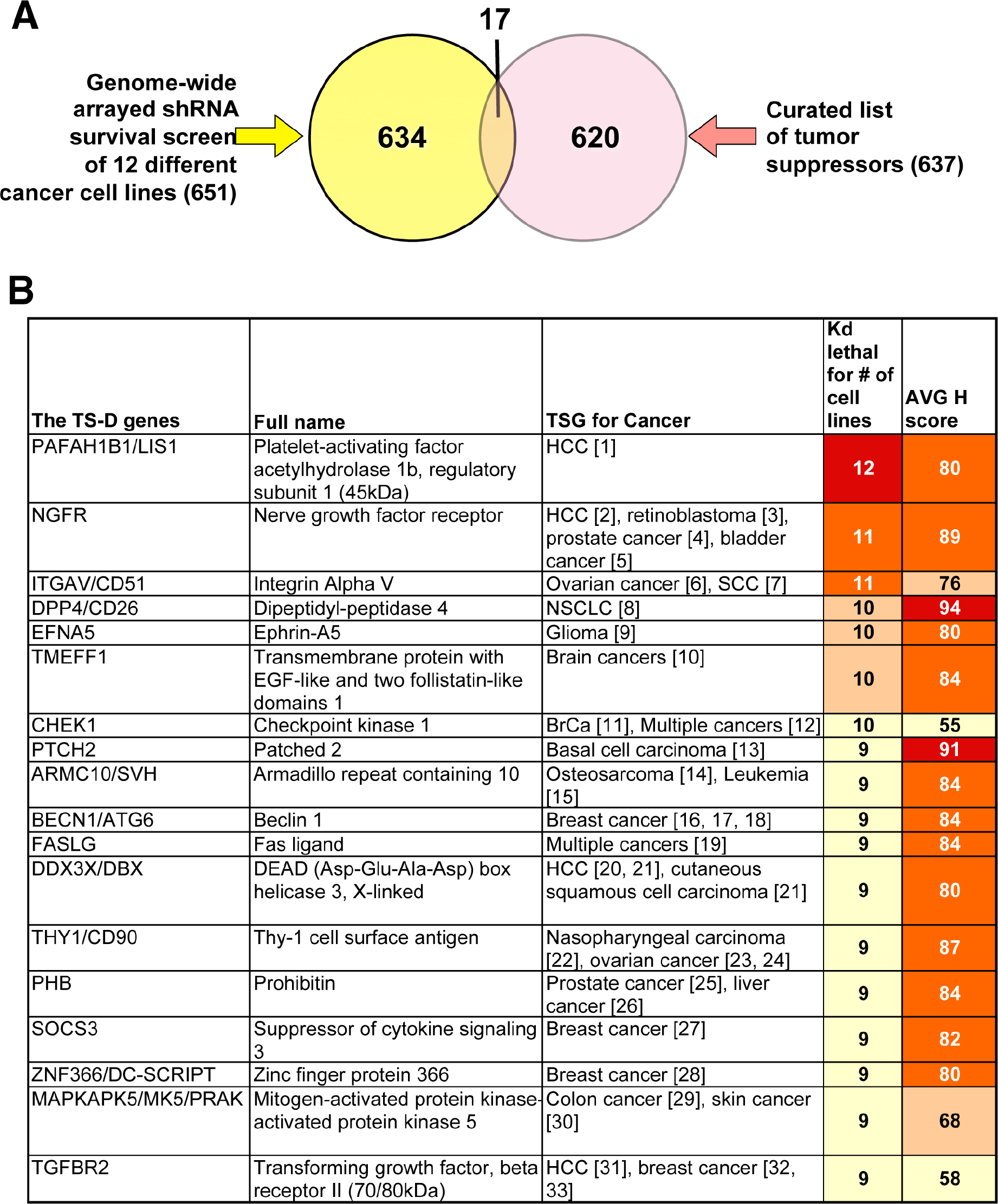
Identification of TS genes among 651 survival genes. (A) Venn diagram showing the overlap of the 651 putative survival genes we identified in 12 genome-wide shRNAs screens with a list of 637 putative tumor suppressors (http://bioinfo.mc.vanderbilt.edu/TSGene). (B) A list of the 17 genes that are putative tumor suppressors and were identified in our lethality screen. The genes are ranked first according to the number of lethality screens in which these genes were found to be survival genes, and second according to the average H score. Higher counts are indicated by darker colors. HCC, hepatocellular carcinoma. FASLG is also shown for comparison. [1], ^1^; [2], ^2^; [3], ^3^; [4], ^4^; [5], ^5^; [6], ^6^; [7], ^7^; [8], ^8^; [9], ^9^; [10], ^10^; [11], ^11^; [12], ^12^; [13], ^13^; [14], ^14^; [15], ^15^; [16], ^16^; [17], ^17^; [18], ^18^; [19], ^19^; [20], ^20^; [21], ^21^; [22], ^22^; [23], ^23^; [24], ^24^; [25], ^25^; [26], ^26^; [27], ^27^; [28], ^28^; [29], ^29^; [30], ^30^; [31], ^31^; [32], ^32^; [33], ^33^. http://bioinfo.mc.vanderbilt.edu/TSGene ^34^.

**Figure S2.**
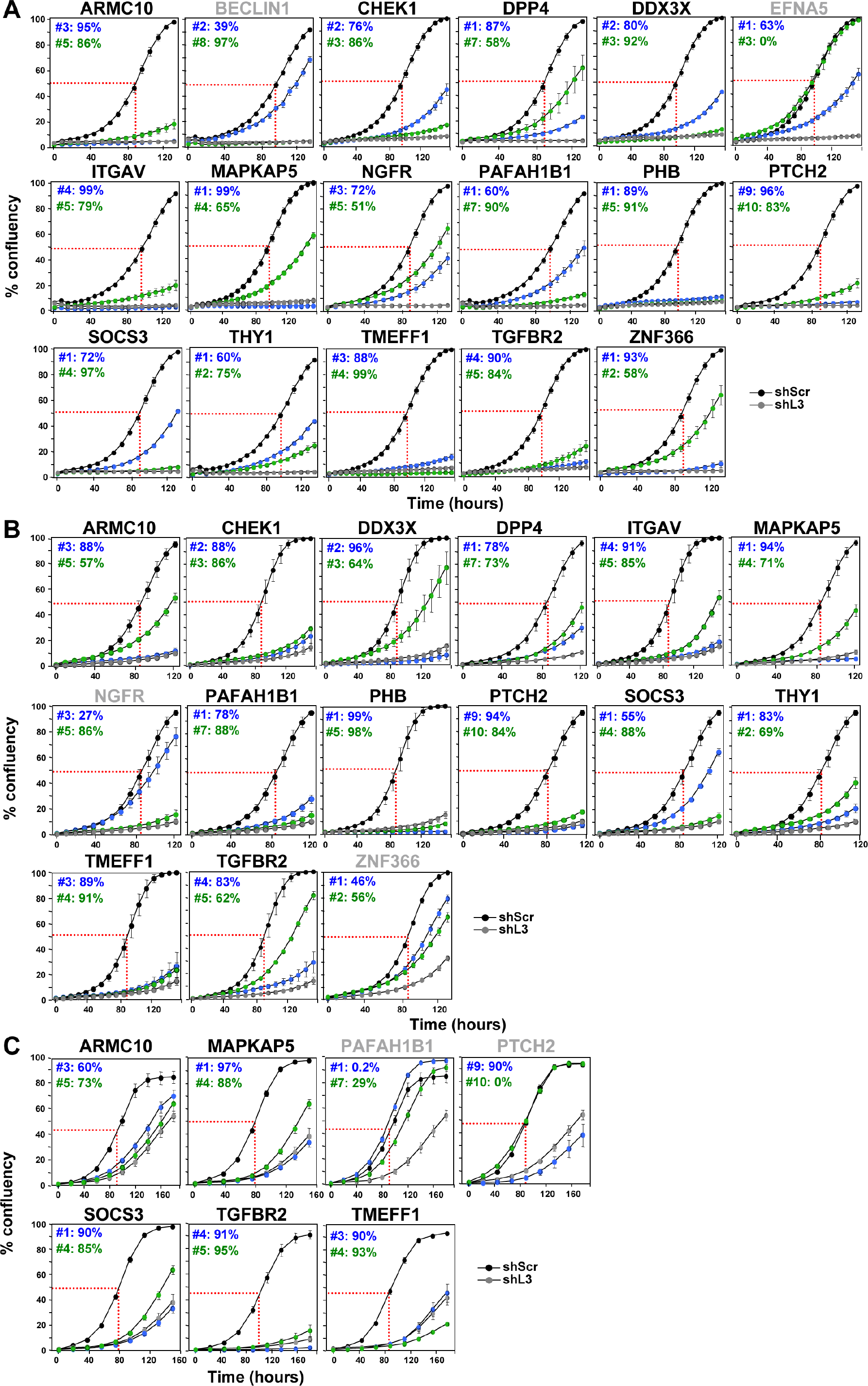
Toxic shRNAs cause growth reduction in T98G, HCT116 and Hap1 cells. Percent cell confluency over time of T98G (A), HCT116 (B) and Hap1 (C) cells infected with shScr, shL3, and two shRNAs derived from each TS gene. The curves for cells infected with two independent shRNA for each TS gene and their specific ID number and respective growth reduction caused by each shRNA are shown in blue and green. Percent growth reduction values (as shown in Table 1) were calculated using STATA1C software when cells infected with shScr reached half maximal confluency as indicated by the red dotted line. Names of genes for which only one of the two shRNAs reduced growth more than 50% are shown in grey.

